# Mechanical Metric for Skeletal Biomechanics Derived from Spectral Analysis of Stiffness matrix

**DOI:** 10.1101/2021.04.29.441973

**Authors:** Petr Henyš, Michal Kuchař, Petr Hájek, Niels Hammer

## Abstract

A new metric for the quantitative and qualitative evaluation of bone stiffness is introduced. It is based on the spectral decomposition of stiffness matrix computed with finite element method. The here proposed metric is defined as an amplitude rescaled eigenvalues of stiffness matrix. The metric contains unique information on the principal stiffness of bone and reflects both bone shape and material properties. The metric was compared with anthropometrical measures and was tested for sex sensitivity on pelvis bone. Further, the smallest stiffness of pelvis was computed under a certain loading condition and analyzed with respect to sex and direction. The metric complements anthropometrical measures and provides a unique information about the smallest bone stiffness independent from the loading configuration and can be easily computed by state-of-the-art subject specified finite element algorithms.

## Introduction

The stiffness is usually defined as a resistance to deformation. This is a natural mechanical behavior and it describes the capacity of a structure to retain an amount of deformation energy. The stiffness should be considered as a combination of both material and structural properties, which form a mechanical response to a given load. The stiffness is a one of the key measures in bone mechanics. A correlation between stiffness and bone strength was shown by several authors^1–4^. The fracture risk was shown to be correlated with bone stiffness^5–9^. The stiffness is also a key component for analyzing the skeletal or intraskeletal adaptation and changes in response to a given loading^1, 10^. The stiffness may also help to better understand the dynamics of the human pelvis during side falls^11^ or alike injury. Moreover, age and sex dependence of stiffness was shown^12–14^: osteoporosis was investigated by analyzing the stiffness^5, 15^.

The bone quality directly influences the patient’s health (i.e., osteoporosis) and hence novel methods for predicting bone mechanics *in-vivo*, ideally non-invasively, are of high interest in contemporary trauma, orthopedics and endocrinology research.

Digital models are often used in biomechanics community to analyze bone mechanics virtually. Nevertheless, such computer models should be driven by clinical patient’s data according to a paradigm of so-called *digital twin*. Although the paradigm of digital twin provides a step towards real clinical applications, the bone mechanics must be precisely known before make fully use of digital models.

Bone stiffness is generally location and direction-dependent, and requires a precise and often complex experimental protocols^11, 13, 16, 17^, or computational approaches such as finite element (FE) models^12, 18–20^. In this study, we introduce an alternative way to estimate bone stiffness. The key idea is to apply spectral decomposition of a stiffness matrix (usually obtained from a FE model) based on the previous authors’ study^21^. The resultant eigenpairs composed from eigenvalues and associated eigenvectors contain a unique information about the modal as well as static stiffness. These pairs have following properties:

A. rotation/translation invariant,
B. naturally considering the structural/intrinsic bone properties (i.e., bone shape, internal structure, bone material, bone elasticity, bone density),
C. under certain conditions provide the minimal bone stiffness for given boundary conditions together with associated deformation shape,
D. does not necessarily require boundary conditions defined.

A) This property allows to compare arbitrarily rotated bone geometries, and it has furthermore been demonstrated in shape analysis^22, 23^. B) The stiffness is constructed often by FE method, where the geometry and material properties are discretized together and represented by a big sparse matrix. C) The stiffness matrix should be symmetric (positive definite) and constant to obtain minimum static stiffness and corresponding eigenvector. D) If no kinematic boundary conditions are provided, the spectrum of eigenvalues close to zero contains the eigenvalues corresponding to rigid movement of the body (it means the three translational and three rotational moves of a body in 3D.) followed by regular deformable eigenvalues. The rigid eigenvalues will not be in the focus of this study.

In this study, we stated a hypothesis that newly formulated mechanical metrics in^21^ provide high sex classification accuracy. Moreover, we compared them with common anthropometric measures and natural frequencies of pelvic bone. We further analyzed the smallest static stiffness of the pelvis with respect to the applied boundary conditions. Our new metric can be considered as modal stiffness, which allows to reconstruct the static stiffness of a bone. Through the study, modal stiffness, smallest static stiffness and natural frequency are called eigenmetric.

### On the Potential Usage of Eigenmetric

It has been shown that bone robustness (the ratio of stiffness and weight) reflects the evolution of skeletal system and how the skeletal system maintains its mechanical function under different enviromental and genetical factors^1, 10, 24^. The robustness is predominantly analysed for long bones, for which the bending stiffness is considered as most relevant^2, 8, 25, 26^ and, according to our approach, the bending stiffness is the smallest one possible for long bones^21^. Moreover the stiffness of long bone can be uniquely expressed for almost any crossection. Nevertheless, it is not clear or it is difficult to define the stiffness for geometrically complex bones or bone assemblies (such pelvic bone or foot^24^ or hip-spine complex). At this point, our eigenmetric comes into play, because it provides the direction (qualitative measure) and magnitude (quantitative measure) of stiffness based only on bone morphology and bone material properties. The eigenvector in our study can be seen as a resultant deformation after aplying certain bone load, which produces a corresponding modal/static stiffness-this raises the question whether loading modes of pelvic bones within the range of physiological loads corresponded to some eigenvectors. Analyzing such load correspondence could help to better understand anthropological or anatomical variations/evolution of bone, for example by proving that bone is physiologically loaded under its static smallest stiffness. The eigenmetric can be also be benefitial in analysis of hip-spine syndrome and load disbalance after hip total replacement or spinal fusion^27–30^. Another potential use of our eigenmetric can be seen in forensic biomechanics as unique bone descriptor (with exaggeration, this given eigenmetric could be called “mechanical DNA of bone”).

## Results

### Modal Stiffnesses and Eigenvectors Description

The first eigenvector for model FEMP-I might represent a torsion deformation with maximal value located in proximity to the pubic tubercle and minimal value below the termination of the anterior gluteal line, see Figure 1. The second eigenvector contains two significant deformation zones and is rather of bending character. Its maximum is located at the iliac crest and minimum at the central part of gluteal surface. The third eigenvector is of a rather complex bending deformation, with tree localizations at different regions of the iliac wing and ischiopubic ramus. The minimum and maximum values are localized close to the anterior superior iliac spine and the dorsal portion of the acetabular margin, respectively. The rest of eigenvectors quickly becomes much more complex and there are no significant deformations interpretable in terms of bending, torsion nor tension. For model FEMP-II, the maximum value for the first eigenvector is located at the anterior portion of the iliac crest. The maximum value for the second eigenvector is projected to ischial tuberosity and the maximum for the third eigenvector represents anterior superior iliac spine. Minimal values of all foregoing modes are the points of fixation, i.e., the symphyseal and auricular surfaces of the pelvic bone. The smallest static stiffness was found with the first modal stiffness and is different for female and male. For female, the mean static stiffness is 170N/mm with standard deviation 48 N/mm for model FEMP-I and mean 97 N/mm with standard deviation 35 N/mm for model FEMP-II. For male, the mean values are 267/206 N/mm with standard deviation 64/59 N/mm for FEMP-I and FEMP-II respective.

**Figure 1.**
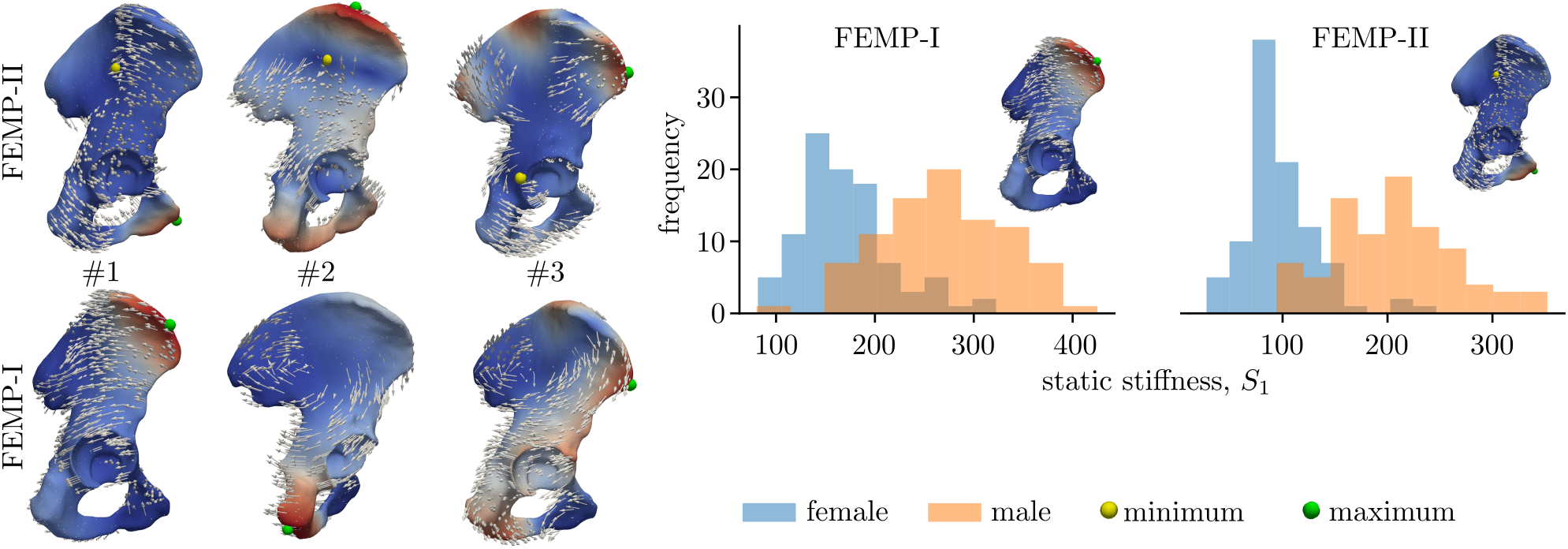
The description of eigenvectors associated with three modal stiffnesses. The arrows represent the direction of deformation. The scalar field represents the magnitude of eigenvector. Minimum markers are not shown for fixed model FEMP-I, given the minima are localized at fixed boundary locations shown in Figure 6. The histograms show the static stiffness [N/mm], based on the first modal stiffness and considered as the smallest static stiffness.

### Comparison of Stiffness Metrics with Anthropometric Measures

#### Fixed Boundary Conditions model FEMP-I

The sex/side classification with modal stiffness in case of fixed model FEMP-I are shown in Figure 2. Only the first modal stiffness allows to classify sex with an accuracy 85%, while none of modal stiffnesses have the potential to classify the side. The other predefined metrics’ classification ability such static stiffness *S* and natural frequency *f* are given in Table 1. The best sex classification accuracy of static stiffness metric *S* as well as natural frequency *f* is reached for first eigenpair. The best accuracy 0.62 for side classification was reached by static stiffness metric *S* for the first eigenpair. In Figure 3 a relation between anthropometric distances defined in^31^ and modal stiffnesses are shown. Only moderate correlations were observed for both sexes with the maximal positive value 0.49 with (CI95%[0.26, 0.67] and *p*\* = 0.0004) for a pair 7-M_9_ for male.

**Figure 2.**
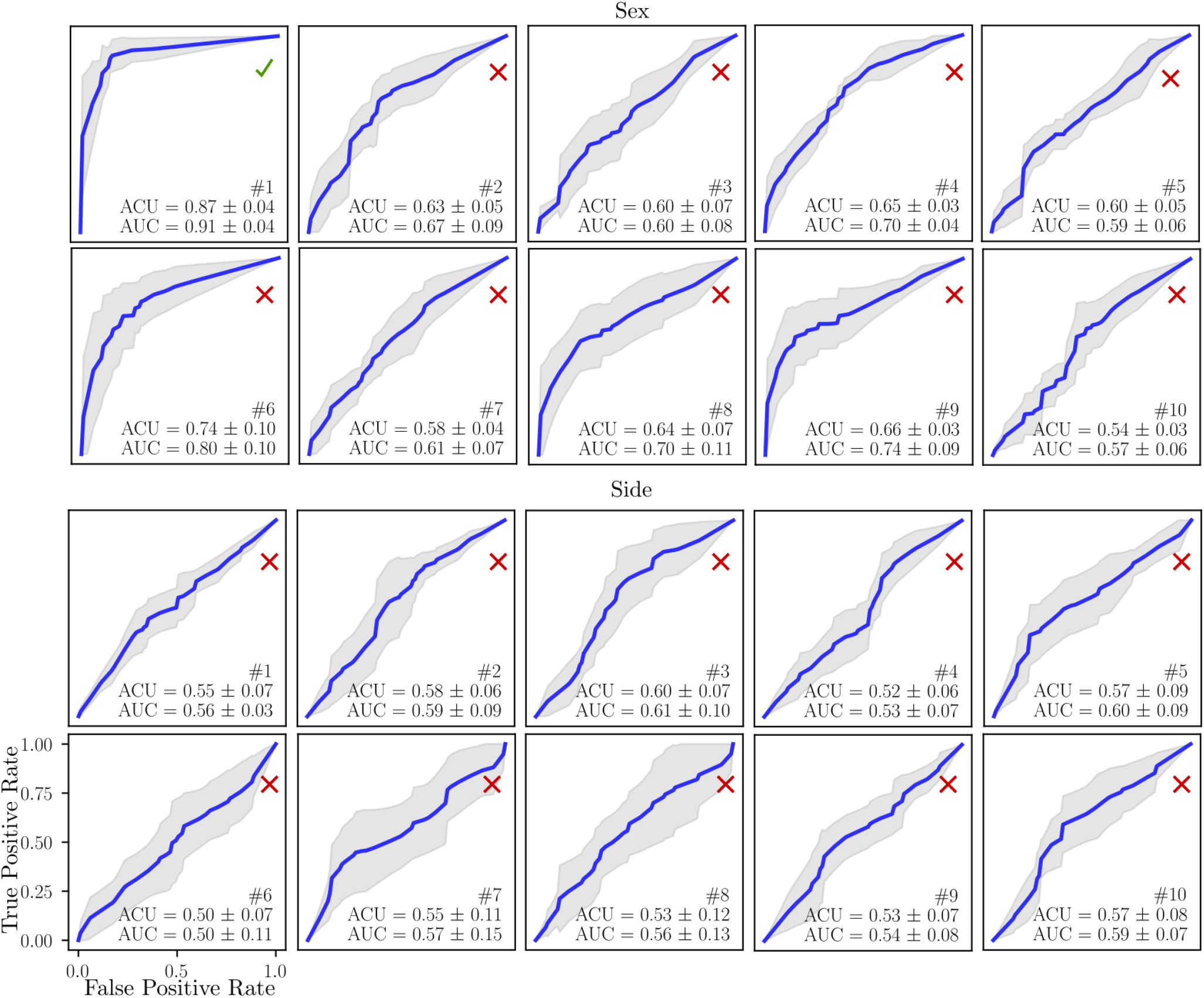
The sex and side classification ability of ten modal stiffnesses *s* for model FEMP-I. The blue curves indicate mean values ROC, grey fill the standard deviation. The mean ACU/AUC with standard deviation is computed within cross-validation step.

**Figure 3.**
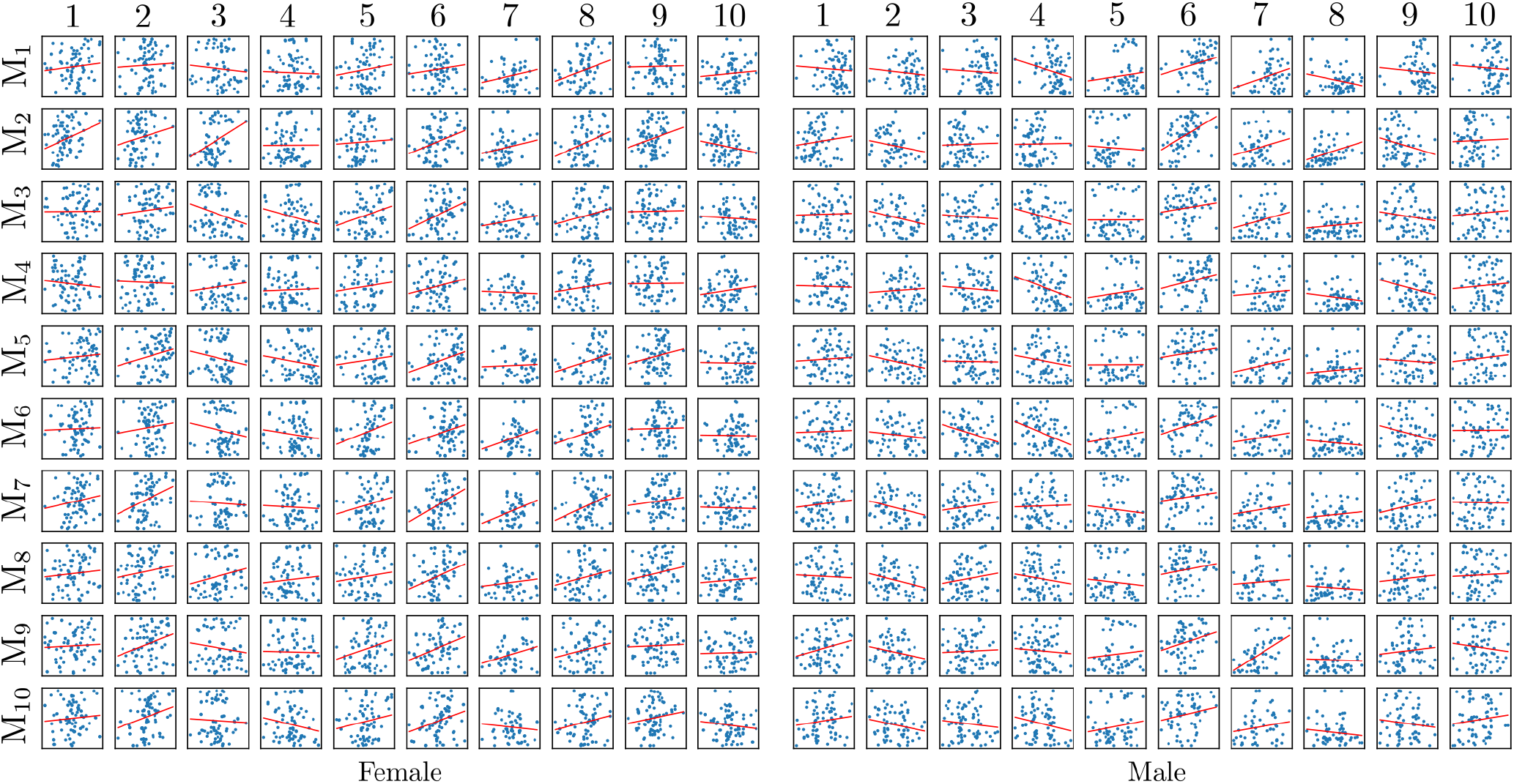
Relation of anthropometric distances and modal stiffnesses computed with model FEMP-I. Blue points represent raw data, while the red curve is regression line fitted.

**Table 1.** Sex and side classification accuracy ACU/AUC of different metrics for model FEMP-I. The CI95% are not shown due to readability, but generally are in range 0.05–0.17

| Metric/eigen number # | 1 | 2 | 3 | 4 | 5 | 6 | 7 | 8 | 9 | 10 |
| --- | --- | --- | --- | --- | --- | --- | --- | --- | --- | --- |
| Sex |  |  |  |  |  |  |  |  |  |  |
| $S$ | 0.82/0.85 | 0.60/0.53 | 0.68/0.66 | 0.67/0.67 | 0.62/0.55 | 0.78/0.79 | 0.70/0.71 | 0.75/0.69 | 0.72/0.73 | 0.66/0.55 |
| $f$ | 0.63/0.58 | 0.63/0.59 | 0.60/0.59 | 0.53/0.54 | 0.53/0.52 | 0.55/0.51 | 0.60/0.58 | 0.54/0.51 | 0.60/0.58 | 0.57/0.54 |
| Side |  |  |  |  |  |  |  |  |  |  |
| $S$ | 0.62/0.53 | 0.58/0.55 | 0.60/0.54 | 0.53/0.52 | 0.49/0.51 | 0.48/0.52 | 0.51/0.51 | 0.52/0.51 | 0.52/0.53 | 0.50/0.51 |
| $f$ | 0.53/0.52 | 0.58/0.55 | 0.55/0.53 | 0.61/0.59 | 0.52/0.51 | 0.55/0.51 | 0.52/0.51 | 0.53/0.52 | 0.59/0.55 | 0.61/0.58 |

#### Free Boundary Conditions Configuration FEMP-II

Sex/side classification accuracy of FEMP-II model is shown in Figure 4. The modal stiffnesses *s*_1_, *s*_5_, *s*_8_ were able to reach a classification accuracy higher than 85%, while none of the modal stiffnesses can classify side. The static stiffness *S*_1_ is also able to predict sex with accuracy higher than 85% according to Table 2. A correlation between modal stiffnesses and anthropometric distances for model FEMP-II is shown in Figure 5. The highest correlation of value 0.55 with CI95%[0.35, 0.70] and (*p*\* = 0.00001) was found for a pair 1-M_2_ for male.

**Figure 4.**
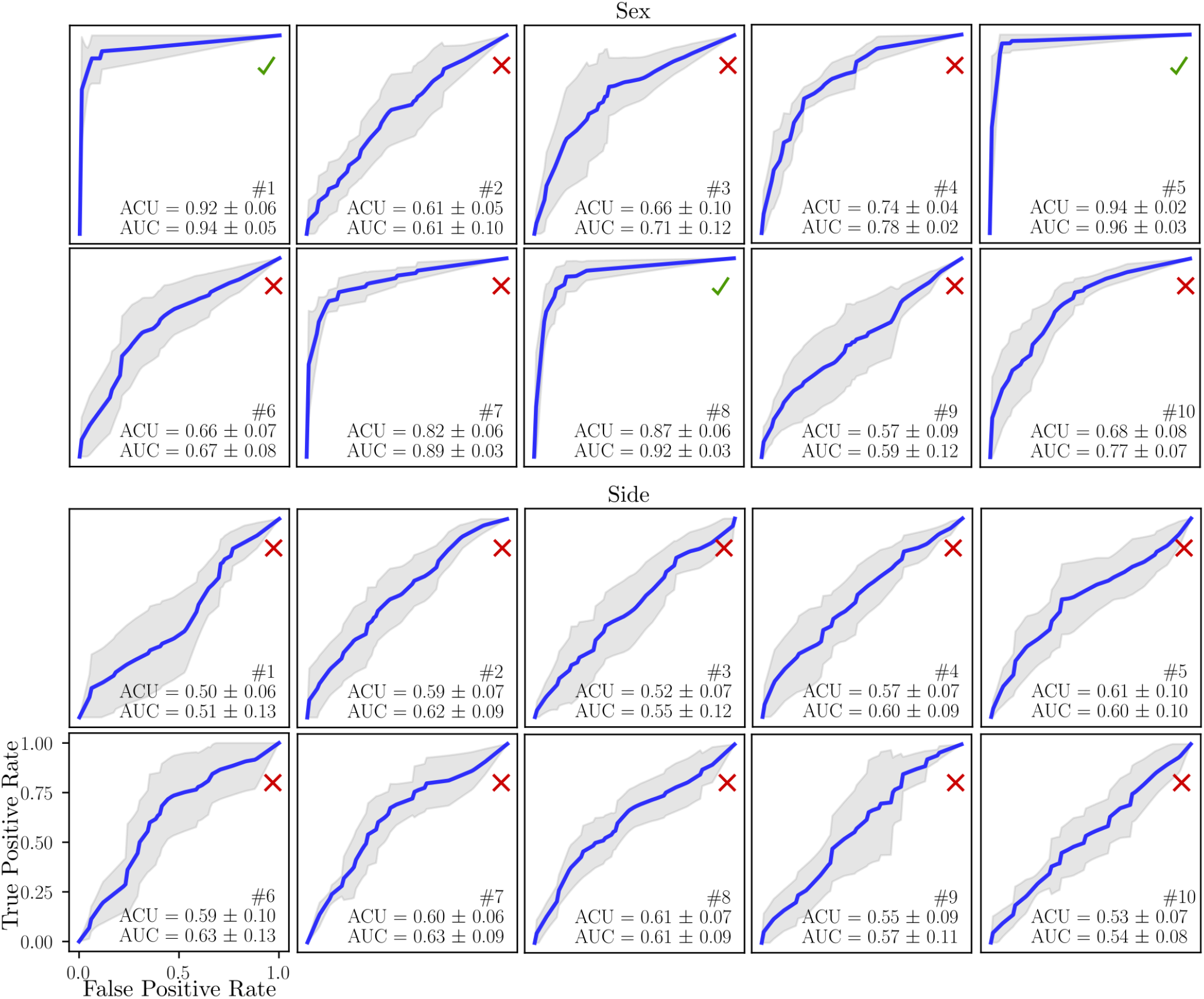
The sex and side classification ability of ten modal stiffnesses for model FEMP-II. The blue curve indicates the mean ROC, while grey fill the standard deviation. The mean ACU/AUC with a standard deviation is computed within cross-validation step.

**Figure 5.**
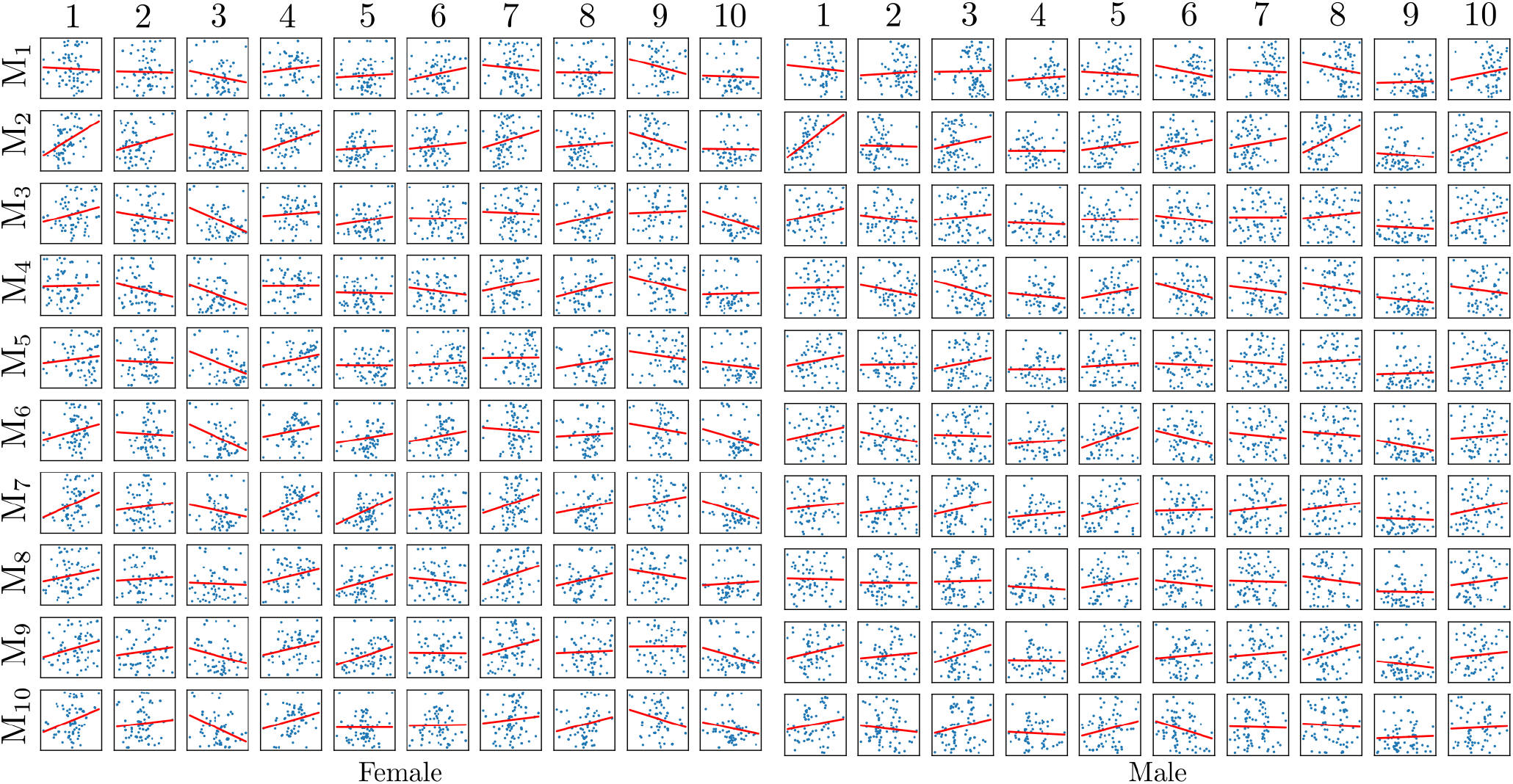
Relation of anthropometric distances and modal stiffnesses computed with model FEMP-II. Blue points represent raw data, while the red curve is regression line fitted.

**Table 2.** Sex and side classification accuracy ACU/AUC of different metrics for model FEMP-II. The CI95% are not shown due to readability, but generally are in range 0.03–0.15

| Metric/eigenpair # | 1 | 2 | 3 | 4 | 5 | 6 | 7 | 8 | 9 | 10 |
| --- | --- | --- | --- | --- | --- | --- | --- | --- | --- | --- |
| Sex |  |  |  |  |  |  |  |  |  |  |
| $S$ | **0.88**/0.89 | 0.65/0.63 | 0.64/0.65 | 0.78/0.79 | 0.75/0.74 | 0.84/0.81 | 0.66/0.67 | 0.84/0.84 | 0.62/0.65 | 0.66/0.55 |
| $f$ | 0.83/0.84 | 0.59/0.61 | 0.59/0.54 | 0.62/0.57 | 0.62/0.54 | 0.60/0.63 | 0.52/0.59 | 0.62/0.61 | 0.55/0.52 | 0.60/0.52 |
| Side |  |  |  |  |  |  |  |  |  |  |
| $S$ | 0.59/0.61 | 0.57/0.55 | 0.52/0.59 | 0.55/0.54 | 0.55/0.61 | 0.59/0.55 | 0.50/0.55 | 0.51/0.54 | 0.44/0.52 | 0.55/0.58 |
| $f$ | 0.55/0.61 | 0.58/0.57 | 0.58/0.62 | 0.59/0.56 | 0.62/0.54 | 0.59/0.61 | 0.59/0.53 | 0.53/0.54 | 0.50/0.53 | 0.56/0.55 |

#### Classification Accuracy of Antropometric Distances

The sex/side classification ability of anthropometric measures M is shown in Table 3. The measures M_2_, M_5_ and *M*_10_ reached classification accuracy at least 85%.

**Table 3.** Sex and side classification accuracy ACU/AUC of antropometric measures M. The CI95% are not shown to improve readability, but generally range within 0.03–0.18

| # | $M_1$ | $M_2$ | $M_3$ | $M_4$ | $M_5$ | $M_6$ | $M_7$ | $M_8$ | $M_9$ | $M_{10}$ |
| --- | --- | --- | --- | --- | --- | --- | --- | --- | --- | --- |
| Sex | 0.59/0.55 | **0.85/0.84** | 0.82/0.83 | 0.70/0.65 | **0.93/0.92** | 0.60/0.55 | 0.79/0.78 | 0.63/0.65 | 0.71/0.72 | **0.85/0.91** |
| Side | 0.54/0.61 | 0.64/0.52 | 0.57/0.55 | 0.44/0.51 | 0.59/0.57 | 0.56/0.59 | 0.39/0.51 | 0.58/0.55 | 0.58/0.54 | 0.54/0.55 |

## Discussion

The mechanical properties of bone are given by the geometry and the intrinsic structure together with material properties. This mix of unique bone properties is highly individual and hence its precise capturing seems a paramount of current *in-silico* biomedical engineering. In this study we provide a unique signature which captures all mentioned bone properties within a set of so-called eigenpairs. Those eigenpairs are derived from a truncated spectral decomposition of bone stiffness and mass represented by algebraic matrices produced by a finite element discretization. Using all eigenpairs computed one could get the full information on bone stiffness (as the stiffness matrix is fully recovered). Practically, only a few eigenpairs are required to capture most of the information about stiffness. In this study, ten modal stiffnesses close to zero were presenting 94% of the bone static stiffness. The smallest eigenvalue corresponded to the smallest natural frequency *f*. Nevertheless, modal stiffnesses do not necessarily follow the same order as natural frequencies since they are rescaled with respect to the power of the eigenvectors amplitude^21^. Interestingly, the modal stiffness corresponding to a first eigenvalue was always the smallest one in our analyses presented here. The corresponding eigenvector maximum magnitude can point to a location and direction of the smallest static stiffness, which can be considered as the most interesting and important. And last, deeper analysis is required to analyze nor explore relations between modal stiffness/smallest static stiffness and important mechanical properties such bone strength, density and anisotropy.

### Interpretation of eigenvectors with respect to static stiffness

The smallest static stiffness can be approximated by modal stiffness if there exist a load and kinematic boundary conditions that produce a deformation of bone, which is similar to an eigenvector. This relation was already demonstrated in the authors’ previous study^21^ and in^32^. This relation can be also explained by a so-called modal reduction technique, where the quantities of interest are projected on to a reduced space spanned by a few eigenvectors to decrease computational complexity while maintaining model as accurate as possible^21^.

### Eigenmetrics predict sex, although are only moderately correlated with anthropometric measures

This fact corresponded to an origin of eigenmetrics, which can be seen as spectral components of bone shape and material information. In fact, the eigenmetrics include apart from geometry information the topological and intrinsic characteristics of bone, which cannot be simply captured by the gold-standard method. Further, we include a side classification as a complementary test to validate tested metrics. As it was expected, no significant side difference was observed from classification tests with tested metrics. The sex classification accuracy of defined metrics comparing to gold-standard seems considerable for individual eigenvalues and depends on the boundary conditions defined. Nevertheless, including used eigenvalues simultaneously into classification algorithm, the final accuracy is excellent (ACC/ACU 98.1%/97.1% ± 0.01%). Moreover, the sex classification accuracy of gold-standard anthropometric landmarks was also comparably high (ACC/ACU 93.3%/92.1% ± 0.01%).

### Boundary conditions affect metrics sensitivity

We demonstrated that natural frequencies, modal and static stiffnesses (eigenmetrics) can be computed for a configuration without boundary conditions defined (FEMP-II). This presents a serious advantage, given the proper modelling of often complicated anatomical boundary conditions is difficult and introduces an additional uncertainty into the model. Moreover, results showed that our metrics computed on the model FEMP-II have better description and classification accuracy than a constrained model FEMP-I. In a sex classification, the FEMP-II metrics contain four sensitive eigenvalues (Figure 4), while FEMP-I metrics only one (Figure 2). Moreover, FEMP-II model can be well validated by an experimental modal analysis with good results^33^. Apart from these advantages, the drawback of FEMP-II model is the difficulty in interpreting the static stiffness as it is computed within an ill-posed static problem. One interpretation could be in an analogy with an alternative model with suitable boundary conditions defined that produces a deformation corresponding to FEMP-II eigenvector. This hypothesis was partially touched in previous author’s study on beam bending analysis^21^.

### Relation of eigenmetrics and modal analysis

The static stiffness is related to modal stiffness in following way. The static stiffness is computed as an inverse sum of modal compliances (compliance is an inverse of stiffness), see relation (8). The modal properties of bone were analyzed to assess the natural frequencies and associated vibration eigenvectors in line with previous experiments^34, 35^ and as a reliable experimental protocol to calibrate computational models^33, 36^. In fact, the squared angular frequencies computed from modal analyses corresponded with the eigenvalues in this study up to scale constant given by modal mass contribution (the mass matrix can be seen as a scaler of eigenvalues, often called in literature as a mass-norm^37^). The eigenvectors are, up to a constant of the same shape as shape from modal analysis. The natural frequencies were considered as a metric as well, which complements modal and static stiffness metrics. Further, our computed natural frequencies of pelvic bones in the dataset are consistent with those in literature^33, 34, 36^.

### Clinical relevance to hip-spine syndrome

The current clinical experience indicates that prior surgical treatment of lumbar spine decreases the outcome of total hip replacement^27, 28^. This unwanted interaction could be explained through lumbosacral-pelvis junction, which can be considered as load-sharing mechanical node^29, 30^. In other words, lumbosacral-pelvis complex could be responsible for load balance in hip and lumbar spine. Therefore, what happens to load balance after total hip replacement or lumbar spine fusion? The overlly stiff hip implants (both femoral stem and acetabular cup) “reinforce” femoral and pelvic bones. Fusing two vertbraes or inserting overly stiff cage between them lead again to increased stiffness (or lower flexibility). Such artificial stiffness may cause a disbalance of the load distribution and hence overloading of other hip-spine regions. This implies implants should be designed such they minimize the load disbalance. Nevertheless, the implant desing strategy is unclear and our stiffness metric could be a good starting point as it provides quantitative (direction) and qualitative (magnitude) of minimall stiffness of bone or bone-implant assembly. Therefore, the influence of implant stiffness on load balance of hip-spine complex is an important topic for next study.

### Study limitations

The presented material model of bone has been defined based on the density–elasticity relation from literature with internal calibration^38^ of used CT. The lack of individual calibration of density–elasticity relationship may potentially decrease the accuracy and reliability of the finite element models used. This potential source of bias will be analyzed in future experiments. Further, although the proposed metrics can be used without boundary conditions defined, we demonstrated that these metrics’ changes are sensitive to boundary conditions. The boundary conditions in this study roughly mimic physiological conditions (model FEMP-I) and their precise definition (potentially including the ligaments and other soft tissues) must be included to extend the usability our proposed metrics. We have obtained that smallest modal stiffness corresponds to a smallest static stiffness. Nevertheless, this observation is rather heuristic as we were not able to provide a suitable prove that the first eigenvector always points to a smallest stiffness, see the study^21^. On the other hand, this limitation does not decrease a potential of proposed metric for studying bone mechanics, because the smallest stiffness can be still found by careful analyzing of stiffness spectra or optimization-based approach proposed in^21^. The smallest static stiffness and corresponding load configuration of a bone have been yet demonstrated on digital models only and will be experimentally verified in a future study. In this study, we simplified material model of bone to be linear and isotropic. This simplification together with density–elasticity relationship could introduce potential sources of errors, which should be carefully investigated in future study. The bone (cortical and trabecular) contains a microstructure that is the source of anisotropy in mechanical properties and significantly alters elastic response and hence the stiffness as well^39–41^. The direction of anisotropy is known for long bones and for mandible^42, 43^. Nevertheles, little is known for the pelvic bone. The artifitial direction^43^ or X-ray based direction field^40, 44^ was considered for mandible bone with good accuracy in mechanical response. These approachech could be the good initial attemp for pelvic bone. Of course, there is also a question how the smallest stiffness is sensitivite to material anisotropy. Therefore, the sensitivity of developed eigenmetrics with respect to material anistropy should be carefully investigated.

## Methods

### Finite Element Model Pipeline

The heterogeneous sample population CT data obtained from anonymized routine CT scans (mean age 64 ± 13.5, gender balanced 100 females/males) consisting of *N* = 200 left/right pelvis bones collected from the Faculty Hospital in Hradec Králové under ethical approval 202102IO2P. The CT data was resampled to have a uniform isotropic resolution 0.8 × 0.8 × 0.8 mm. To extract the geometry, the pelvic bones were segmented using a semi-automatic method based on the interactive GraphCut algorithm^45, 46^. The masks from CT segmentation were converted to an STL (stereolitography format) representation (VTK library^47^). Consequently, the volume finite element meshes were automatically built with library TetWild^48^, see Figure 6. The CT values were transformed to an effective density in order to compute total mass of the bones *ρ*_eff_ = *b*CT, where the scaling coefficient is *b* = 658 g/cm^3^^49^. To estimate Young’s modulus, the CT scans were calibrated internally resulting in hydroxyapatite content in bone^38^. The density–Young’s modulus in [MPa] relationship

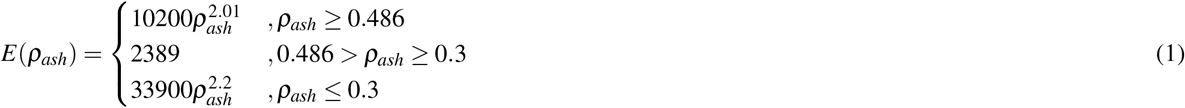

was used where *ρ_ash_* = 0.877*ρ_HA_*+ 0.08 g/cm^3^^50–52^. The Poisson’s ratio *ν* is defined as a constant 0.3^49^ over the whole bone domain. Two sets of boundary conditions were built. The first model FEMP-I described the fixed boundary conditions, see Figure 6. The fixed boundary conditions were inspired by previous works^53^ and mimic the physiological conditions up to some extent. The second model considers no kinematic and no force boundary conditions, called FEMP-II. Based on the bone geometry, isotropic linear model of bone and boundary conditions defined, the stiffness and mass matrices K, M are constructed with help of finite element method in a usual sense:

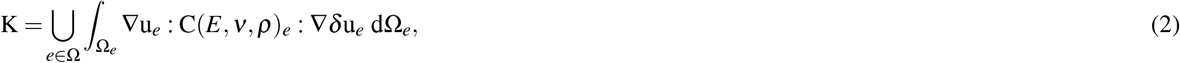

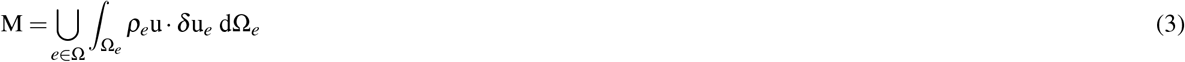

**Figure 6.**
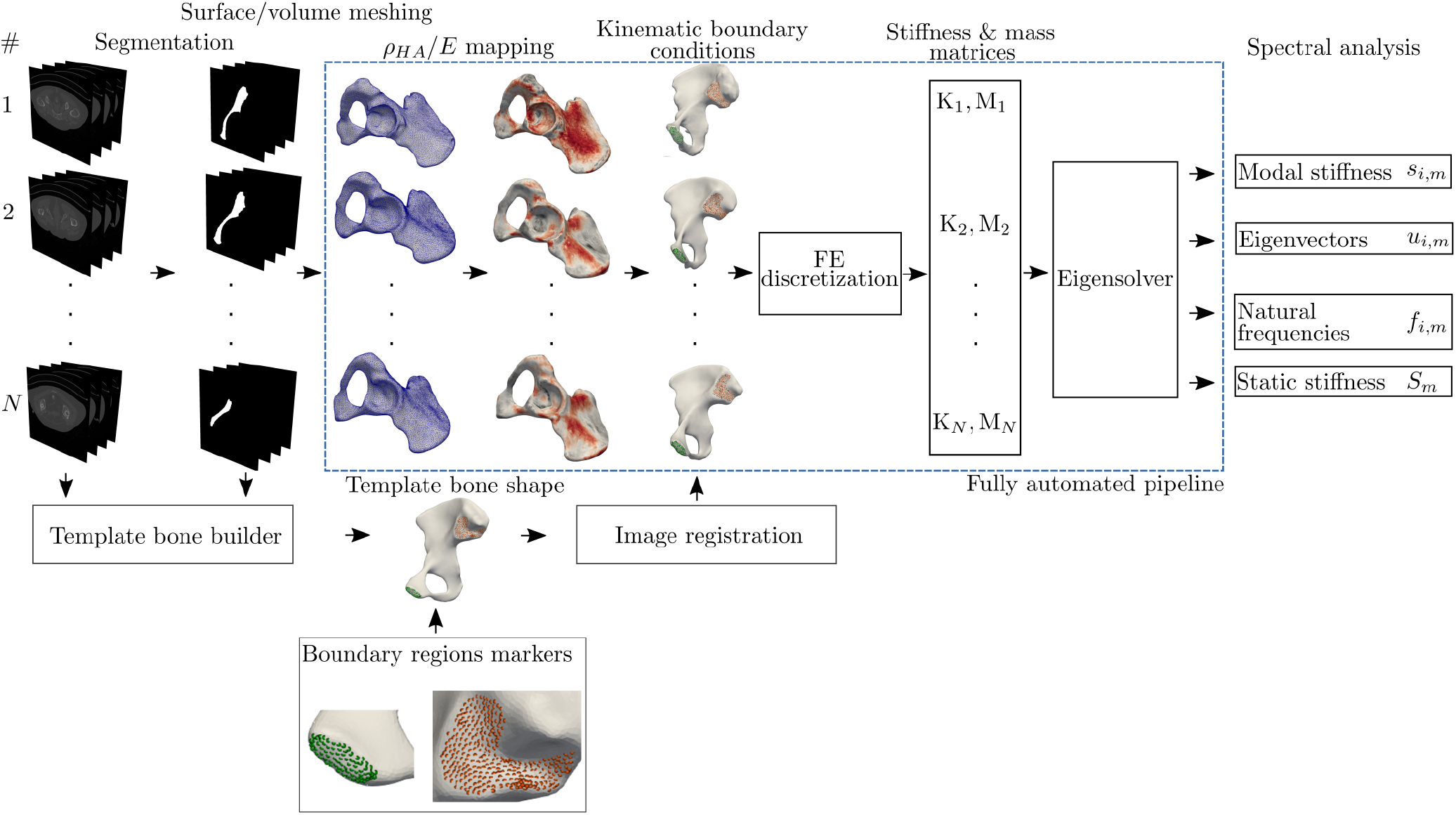
A pipeline of building the FE model based on the CT data. The boundary points(green/orange) are used to apply the kinematic boundary conditions.

The operator ∇ is a symmetric gradient of displacement u. The displacement u and its variation *δ*u are approximated with an arbitrary linear piece-wise continuous functions (in Fenics notation: P1 Lagrange finite element^54^), while C*_e_* represents the tensor of material coefficients inferred from density–elasticity relationship and approximated on the discontinuous finite element space of zero order (i.e., in Fenics notation DG0). The stiffness matrix was discretized on tetrahedral mesh domain Ω representing the bone geometry. The characteristic element size is 1 mm, which corresponds to 700 000–1 000 000 elements. The element size is estimated at auxiliary convergence study on five eigenvalues, which should change up to 5% between two mesh resolutions. Once the element particular quantities are constructed, the global stiffness matrix K is assembled (assembling operator ∪). The details of presented FE discretizations of stiffness matrix can be found for example in^54,55^. The homogeneous fixed boundary conditions are injected into stiffness matrix K on an algebraic level by proper zeroing rows and columns (see, the manual for Fenics library^54^). Once the stiffness matrix is built up, the following generalized eigenvalue problem (modal analysis) can be formulated:

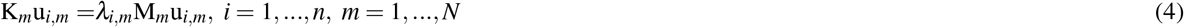

where *λ_i,m_* and vector u*_i,m_* are *i*’th eigenvalue and eigenvector associated with of sample *m*, respectively. In order to compute those eigenpairs, LOBPCG solver from SLEPc library was used to find 10 smallest real eigenpairs^56^. The overall computational framework is written within problem solving environment library Fenics 2018.1^54^.

### Eigenmetrics Definition

The norm of the eigenvector ||*u_i_*|| of a *m*’th sample is defined as a pointwise norm:

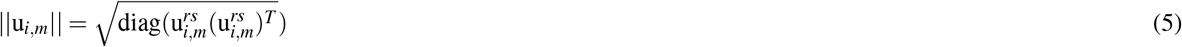

where eigenvector 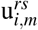 is reshaped to have a new dimension, which contains pointwise three components *x, y, z* of a vector in 3D. To localize the point where the stiffness of structure is potentially lowest one, it is possible to find a maximum of above norm of eigenvector and corresponding index *k* (degrees of freedom in FE model) of point. This corresponds with looking for maximum of compliance:

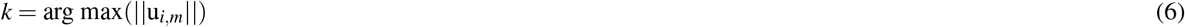

Once we have determined the index *k* we can compute *i*’th modal stiffness of *m*’th sample as

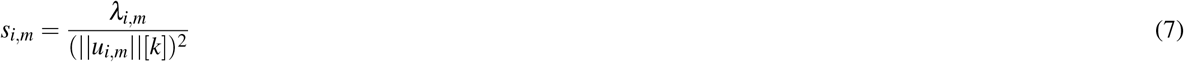

The static stiffness at point with index *k* is computed in virtue of modal superposition described in^21^ and can be expressed as:

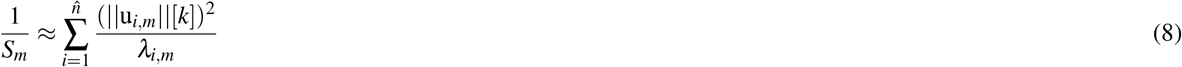

where 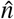 is the number of eigenpairs (eigenvalue, eigenvector) included. Static stiffness can also be understood as the ratio of force and deformation. With ten eigenpairs, approximation error in static stiffness was 4%, which is acceptable in terms of speed and accuracy of truncated spectral decomposition. The natural undamped frequency is defined as usual:

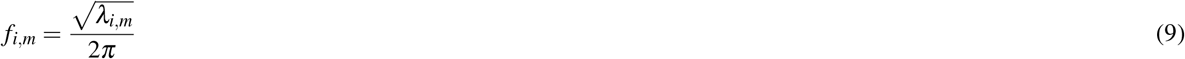

The proposed definitions of modal stiffness *s_i_* as well as static stiffness *S* were based on the theoretical and computational study^21^.

### Anthropometric Analysis of Pelvis

The shape of the pelvic bones were parametrized by a series of landmarks called *B*_1_, *B*_2_,…, *B*_19_ and associated distances M_1_, M_2_,…, M_10_, see author’s previous study^31^. These landmarks and distances have been adopted from DSP2 tool (Diagnose Sexuelle Probabiliste^57^). DSP2 is capable of sexing the bone specimens by computing the probability of being male or female using the combination of sex sensitive variables. Chosen landmarks and distances do not only describe the shape of bone, additionally they provide a set of sex-specific information. Pelvic morphology is complex, sexually dimorphic, and hence is considered as a good bone model for analysis of eigenmetric proposed in this study. Additionaly, the information about the side (left/right pelvis bone) was included in the study.

### Automatized Construction of Boundary Conditions and Anthropometric Landmarks

To decrease the operator error in construction of large number of FE models, the automatized generation of landmarks and boundary conditions was employed with the help of deformable registration algorithm^31^, see Figure 6. The template bone shape was estimated based on^58^ and consequently between the template bone and all bones in datasets the symmetric, diffeomorphism map was computed by Symmetric registration method (SyN) with demon metric (ANTs library^59^). The correlation between automatically and manually computed anthropometric distances is shown in Table 4. The highest correlation was achieved for the first operator and the distance *M*_3_, while the lowest correlation was achieved for the distance *M*_4_ measured by first operator. Moreover, the sex classification trained by automatically computed anthropometric distances performs slightly better in comparing to manually computed distances(ACU/AUC 0.87±0.1/0.96±0.1 with all distances included at once versus ACU/AUC 0.85±0.1/0.93±0.1 with all distances included). The smallest correlation was obtained for fifth modal stiffness *s*_5_ in case of first operator, see Table 5. The sex prediction performance was achieved a similar for both manual and automatic approaches with all modal stiffnesses included at once (ACU/AUC 0.91±0.2/0.94±0.2). To determine the accuracy of the automatic boundary marking, in the Table 5 the correlation metrics are computed.

**Table 4.** Correlation metrics between automatically and manually computed anthropometric distances.

| Distance, M | Operator I: Pearson's.coef |  | Operator I: CI(95%) |  | Operator II: Pearson's.coef |  | Operator II: CI(95%) |  |
| --- | --- | --- | --- | --- | --- | --- | --- | --- |
|  | rep. #1 | rep. #2 | rep. #1 | rep. #2 | rep. #1 | rep. #2 | rep. #1 | rep. #2 |
| 1 | 0.94 | 0.94 | [0.91, 0.97] | [0.90, 0.97] | 0.92 | 0.98 | [0.91, 0.95] | [0.94, 0.99] |
| 2 | 0.94 | 0.94 | [0.91, 0.97] | [0.90, 0.97] | 0.90 | 0.95 | [0.88, 0.93] | [0.97, 0.99] |
| 3 | 0.98 | 0.98 | [0.97, 0.99] | [0.97, 0.99] | 0.95 | 0.93 | [0.93, 0.99] | [0.91, 0.98] |
| 4 | **0.77** | 0.80 | [0.63, 0.87] | [0.67, 0.88] | 0.81 | 0.81 | [0.79, 0.84] | [0.80, 0.85] |
| 5 | 0.84 | 0.99 | [0.75, 0.91] | [0.99, 1.00] | 0.83 | 0.91 | [0.74, 0.91] | [0.85, 0.96] |
| 6 | 0.95 | 0.93 | [0.92, 0.97] | [0.89, 0.96] | 0.95 | 0.93 | [0.92, 0.97] | [0.89, 0.96] |
| 7 | 0.96 | 0.97 | [0.94, 0.98] | [0.96, 0.99] | 0.91 | 0.95 | [0.89, 0.94] | [0.93, 0.98] |
| 8 | 0.84 | 0.77 | [0.73, 0.91] | [0.64, 0.87] | 0.87 | 0.82 | [0.79, 0.92] | [0.75, 0.89] |
| 9 | 0.97 | 0.97 | [0.96, 0.99] | [0.96, 0.99] | 0.95 | 0.96 | [0.92, 0.99] | [0.92, 0.99] |
| 10 | 0.96 | 0.97 | [0.94, 0.98] | [0.95, 0.98] | 0.93 | 0.95 | [0.92, 0.98] | [0.93, 0.98] |

**Table 5.** Correlation metrics between modal stiffness computed on model FEMP-II with automatically and manually marked boundary conditions.

| Modal stiffness, s | Operator I: Pearson's coefficient |  | Operator I: CI(95%) |  | Operator II: Pearson's coefficient |  | Operator II: CI(95%) |  |
| --- | --- | --- | --- | --- | --- | --- | --- | --- |
|  | rep. #1 | rep. #2 | rep. #1 | rep. #2 | rep. #1 | rep. #2 | rep. #1 | rep. #2 |
| 1 | 0.91 | 0.93 | [0.82, 0.96] | [0.89, 0.98] | 0.90 | 0.91 | [0.82, 0.94] | [0.87, 0.98] |
| 2 | 0.90 | 0.93 | [0.85, 0.93] | [0.90, 0.97] | 0.87 | 0.90 | [0.81, 0.91] | [0.85, 0.94] |
| 3 | 0.96 | 0.99 | [0.90, 0.99] | [0.93, 0.99] | 0.95 | 0.91 | [0.90, 0.99] | [0.87, 0.97] |
| 4 | 0.91 | 0.94 | [0.84, 0.96] | [0.90, 0.98] | 0.88 | 0.89 | [0.81, 0.92] | [0.84, 0.93] |
| 5 | **0.73** | 0.81 | [0.66, 0.84] | [0.75, 0.89] | 0.83 | 0.79 | [0.76, 0.83] | [0.72, 0.84] |
| 6 | 0.84 | 0.95 | [0.80, 0.91] | [0.91, 0.98] | 0.90 | 0.91 | [0.85, 0.94] | [0.87, 0.95] |
| 7 | 0.91 | 0.94 | [0.85, 0.96] | [0.90, 0.98] | 0.89 | 0.91 | [0.84, 0.94] | [0.86, 0.96] |
| 8 | 0.87 | 0.93 | [0.79, 0.91] | [0.88, 0.87] | 0.83 | 0.87 | [0.77, 0.89] | [0.79, 0.92] |
| 9 | 0.92 | 0.97 | [0.89, 0.96] | [0.89, 0.99] | 0.85 | 0.91 | [0.80, 0.89] | [0.83, 0.98] |
| 10 | 0.91 | 0.94 | [0.86, 0.98] | [0.87, 0.98] | 0.88 | 0.93 | [0.81, 0.91] | [0.89, 0.98] |

### Statistical Evaluation

Ensemble random forest method was used to binary classify sex and side. The relation between anthropometric variables and eigenmetrics was analyzed with Spearman’s correlation on a significance level 95%. The classification is evaluated with sensitivity/specificity/area under curve (AUC) metrics summarized in receiver operation characteristic (ROC) based on 3-fold cross-validation procedure^60^. The Pearson’s correlation was used to measure the degree of correspondence of automatized and manually measured anthropometric distances M. To achieve a reliability of manual measuring of distances, two operators with two repetitions computed the anthropometric distances of 50 samples, hence four correlation values are reported. The same reliability test was used to analyze manual and automatic computing of boundary conditions markers. If possible, the 95% confidence interval (CI) is computed.

## Acknowledgements

The authors acknowledge support from the German Research Foundation (DFG) and the University of Leipzig within the program of Open Access Publishing.

## Author contributions statement

P.H. prepared theoretical basis, M.K. Anthropometrical analysis, N.H. Anatomical description and manuscript preparation, P.H. thorough manuscript revision. All authors reviewed the manuscript and agreed with its form.

